# Hydrogen sulfide exposure reduces thermal set point in zebrafish

**DOI:** 10.1101/2020.02.06.935957

**Authors:** Dimitri A. Skandalis, Cheryl D. Dobell, Joshua C. Shaw, Glenn J. Tattersall

**Affiliations:** Department of Biological Sciences, Brock University, St. Catharines, ON, Canada L2S 3A1

**Keywords:** thermal preference, set point, thermoregulation

## Abstract

Behavioural flexibility allows ectotherms to exploit the environment to govern their metabolic physiology, including in response to environmental stress. Hydrogen sulfide (H_2_S) is a widespread environmental toxin that can lethally inhibit metabolism. However, H_2_S can also alter behaviour and physiology, including a hypothesised induction of hibernation-like states characterised by downward shifts of the innate thermal setpoint (anapyrexia). Support for this hypothesis has proved controversial because it is difficult to isolate active and passive components of thermoregulation, especially in animals with high resting metabolic heat production. Here, we directly test this hypothesis by leveraging the natural behavioural thermoregulatory drive of fish to move between environments of different temperatures in accordance with their current physiological state and thermal preference. We observed a decrease in adult zebrafish (*Danio rerio*) preferred body temperature with exposure to 0.02% H_2_S, which we interpret as a shift in thermal setpoint. Individuals exhibited consistent differences in shuttling behaviour and preferred temperatures, which were reduced by a constant temperature magnitude during H_2_S exposure. Seeking lower temperatures alleviated H_2_S-induced metabolic stress, as measured by reduced rates of aquatic surface respiration rate. Our findings highlight the interactions between individual variation and sublethal impacts of environmental toxins on behaviour.

## Introduction

Environmental toxicants may act through myriad pathways, including hijacking the body’s own signalling pathways. Hydrogen sulfide (H_2_S) is a widespread aquatic toxicant that is also an important endogenous gasotransmitter, and occurs naturally through anoxic decomposition (e.g., salt marshes and mangrove swamps) or due to anthropogenic activities (e.g., sewage treatment and aquaculture farming, [1,2]). Exogenous H_2_S inhibits aerobic respiration and, together with low oxygen (hypoxia), contributes to large fish kills [1–3]. However, endogenous H_2_S is involved in the response to hypoxia and has physiological regulatory roles in synaptic activity and cognitive function, and inflammation [4–6]. Environmental H_2_S could therefore induce potent physiological responses independently of metabolic distress. For instance, it has been proposed that application of exogenous H_2_S in combination with low temperatures induces a drop in body temperature through entry into a hypometabolic hibernation-like state in mice [7]. However, it is unclear if this is an effect of H_2_S alone or aggravation of a conserved environmental hypoxia response [8,9]. These studies have been performed in small mammals with their thermoneutral zone, where thermogenesis and dissipation are normally balanced; metabolic poisoning by exogenous H_2_S might impair resting heat production rather than stimulate a controlled depression of the set point. In contrast, the thermal setpoint of ectotherms like fish is regulated behaviourally, enabling direct assessment of body temperature regulation. Whereas in most terrestrial animals exogenous H_2_S is applied to study the gasotransmitter’s endogenous functions [7,8], in aquatic habitats exogenous H_2_S is also ecologically relevant [10–13]. We exploit this physiology as a direct test of the hypothesis that H_2_S drives changes in thermal preferences, which is significant for the ecology and behaviour of this major taxon.

Fish select environments based on their preferred temperatures and behavioural motivation. Active fish, like zebrafish (*Danio rerio*), tend to move toward their preferred temperatures, whereas benthic or sedentary species tend to remain in place until extreme temperatures become unbearable [14]. Fish can detect water temperature changes of 0.05°C or less [15]. Preferred temperatures vary within and among individuals, depending on factors such as growth, acclimation, health, and social cues [16–22]. Preferred temperature reflects the fish’s metabolic state [17], and numerous fish select colder temperatures (i.e., behavioural anapyrexia) in hypoxia than in normoxia conditions [23–25]. Presumably, the response is due to enhanced haemoglobin oxygen-binding capacity and reduced metabolic demand of tissue at low temperatures, which balance oxygen supply and demand [23]. Exposure of fish to H_2_S shares many physiological similarities with hypoxia [1,25], which could arise because H_2_S metabolism functions as an endogenous O_2_ sensor [13,25]. Here, we examine how zebrafish thermal preferences are altered with exposure to H_2_S in normoxic conditions. We hypothesized that H_2_S would trigger a reduction in individual thermal setpoint, which would suggest that sublethal H_2_S can also have major effects on physiology and behaviour.

## Materials and Methods

### Animals and husbandry

Zebrafish (*Danio rerio*) from a local supplier were housed in 40L aquarium tanks at 27°C and pH 7.6-7.8 (Seachem^™^ Acid Buffer), on a 12:12 light:dark cycle with once-daily feedings (Tetra Flakes^™^). Fish were housed at least 24 days and habituated to walls lined with white contact paper prior to experiments.

### Shuttlebox design and gas mixing

Thermal preferences were tested in a two-chamber dynamic shuttlebox described previously [26] by automatically tracking body position (x,y; *x*=0 at midline, +*x* to the right), swim velocity, and chamber temperatures; each variable was sampled at 1 Hz and stored for offline analysis by the program ICFish (v.2.1, Brock University Electronics; see [26]). Hydrogen sulfide was bubbled through inaccessible side chambers into the main chamber, allowing gas mixing without disturbing the fish. Air and 0.2% H_2_S were mixed to achieve the appropriate H_2_S concentration (0% or 0.02% H_2_S) by flow meters (Omega rotameters) at 5000 mL min^−1^. We found 0.02% H_2_S elicited robust responses without severe distress found at 0.07% H_2_S (not shown). Bubbling mixed gases avoids difficulties in determining H_2_S concentration from dissolved NaS salts [5,25], and balancing H_2_S with air (rather than nitrogen) guarantees normoxia (20.88% O_2_). Gas dissolution equilibrated for 30 minutes. Gas was also flowed under a clear Plexiglas cover to maintain constant chamber gas pressures, minimise condensation, and preclude gas gradients within the chamber (e.g., lower H_2_S at the water surface).

### Respiratory drive in response to H_2_S

Respiratory responses were assessed in two fish at a time at a constant temperature, each placed in one side of the shuttlebox and separated by an opaque barrier. Six individuals were exposed to each combination of 0 or 0.02% H_2_S and 21 or 28°C and two individuals to 0% H_2_S at 28°C, for 60 min. Body position in the water column was sampled at 1 Hz from a horizontal view, and frames with fish at the surface (successes) out of the total number of frames (trials) were interpreted as the aquatic surface respiration rate ([24], analyzed in ImageJ v1.52). The effects of H_2_S and temperature were modeled with a binomial error distribution using Markov chain Monte Carlo (*brms* [27]) with four chains of 10,000 iterations each, run to convergence (*Ȓ* = 1 [28]), discarding half as burn-in. Significance was interpreted as posterior parameter credible intervals excluding zero.

### Shuttling experiments

Pilot experiments revealed the most robust thermoregulatory behaviour when fish could first learn to associate each chamber with a constant temperature difference. Therefore, fish were introduced to the left chamber (Figure 1, set to 1.5°C below housing temperature) and habituated one hour with a constant 3°C difference between chambers. Ramping then commenced for two hours, triggered when the fish entered the left (cooling, −0.5°C min^−1^) or right (warming, +0.5°C min^−1^) sides, within limits set to 15 and 35°C (within 5°C of *D. rerio*’s thermal tolerances [19]). Following gas equilibration for 30 minutes, behaviour was recorded for four hours (test phase). Fish that exhibited severe distress (e.g., loss of equilibrium) were pre-emptively removed from the experiment.

**Figure 1.**
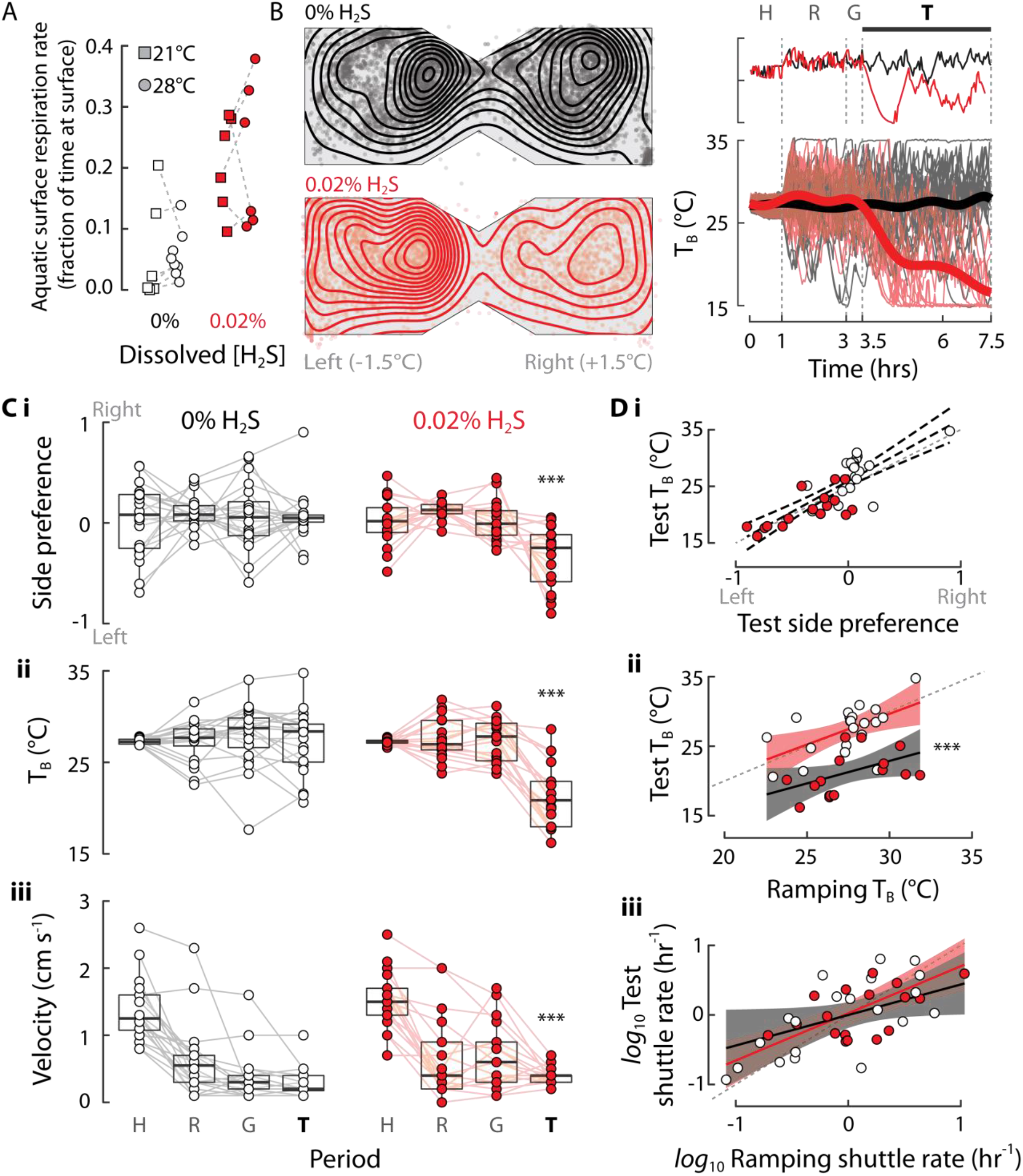
Hydrogen sulfide (H_2_S) exposure drives zebrafish behavioural anapyrexia. **A** Adult zebrafish exposed to 0.02% H_2_S (red) exhibit greater aquatic surface respiration rates (ASR, fraction of time at surface) compared to 0% H_2_S, but this was reduced by lower body temperatures, T_B_ (H_2_S Credible Intervals, CrI: 1.41–4.36; ΔT_B_ CrI: 0.11–2.43). **B** Adult zebrafish actively defend T_B_ by shuttling between chambers of 3°C difference. Zebrafish habituated (H) to constant temperature difference over 1 hr, then temperature ramped (R) up or down according to the selected arm. Following a 30 min gas (G) equilibration phase, testing (T) was performed for four hours. Two-dimensional kernel density plots of average fish positions (not constrained by chamber boundaries) reveal fish prefer the cooler chamber (left) with exogenous 0.02% H_2_S (red). Minor online tracking errors (points outside boxes) are shown for analytical transparency but do not affect results. Average body temperature (right, thick lines) is effectively constant at 0% H_2_S but rapidly decreases in 0.02% H_2_S (thin lines are individual traces). Representative traces from two individuals (above) demonstrate continuing shuttling behaviour in both conditions. **C** Measured responses in each time period (B) show the progression of responses over the course of the experiment, including between ramping and test phases. *i* Significant side preferences were observed only in 0.02% H_2_S (time ratio; 0% H_2_S confidence interval, CI: - 0.06–0.18; 0.02% H_2_S CI: −0.48– −0.21). *ii* Preferred temperature is significantly lower at 0.02% H_2_S (*p*<0.001; CI ΔT_B_: 3.6–8.4°C). *iii* Fish activity, measured by velocity, declined over time at 0% H_2_S, presumably reflecting decreased exploration, but increased with the introduction of H_2_S, including during the test phase (*p*<0.001; CI Δ velocity: 0.21–0.68 cm s^−1^). **D** *i* Interindividual variation in T_B_ across treatments was correlated with time spent in each chamber (*p*=0.001). All individuals must lie on this trend by construction, so treatment effects were excluded (distinguished by dashed CI band). *ii* Preferred T_B_ was repeatable from ramping to test phases (*p*=0.006) with slope near unity (0.88, CI: 0.24–1.53). A significant effect of 0.02% H_2_S without detectable interaction (*p*<0.001; CI ΔT_B_: 4.1–8.3°C; T_B_ ✕ H_2_S: *p*=0.61) points to a shift in T_B_ set point. *iii* Shuttle rate was repeatable among individuals (*p*=0.01) but with slope less than unity (CI: 0.38–0.98), suggesting fish learn to defend T_B_ even while expending less effort (fewer shuttles). The relationship was unaffected by H_2_S (H_2_S *p*=0.85; *log* shuttling rate ✕ H_2_S: *p*=0.37). Sample sizes: *n* 0% H_2_S=20; *n* 0.02% H_2_S=17.

### Data collection

Fish *x* position was used to calculate shuttling rate (frequency of crossing *x*=0, min^−1^) and side preference 2•(0.5-Time_x<0_/Time_total_). Thermal inertia of small fish is minimal compared to water temperature [18,29,30], so we calculated body temperature (T_B_) by averaging current chamber temperature with T_B_ in the previous time step. Lower and upper escape temperatures (LET and UET) were the last recorded T_B_ prior to a shuttle.

Qualitative differences of side preference were visualised through two-dimensional kernel density estimates with a bandwidth of 50 x 50 pixels, unconstrained by shuttlebox boundaries. For visualisation, the shuttlebox walls were estimated *post hoc* from fish positions, and the resulting densities clipped to those borders. We applied generalised additive models (R package *mgcv* [31]) to model the differences in average T_B_ over time between 0% (reference spline) and 0.02% (difference spline) H_2_S. Serial autocorrelation of time series model errors was incorporated through a Gaussian process spline basis with AR(1) autocorrelation structure. All other variables were quantitatively analysed through linear models. Fish velocity and shuttling rate were log-transformed prior to analysis. To quantify individual variation in responses to H_2_S, we assessed the relationships of responses during the ramping phase to those during the testing phase. This approach is predicated on the consistency of intra-individual thermal preferences over the course of the experiment, which we justify by calculating repeatabilities (*rptR* [32]) of thermal preferences and behaviour between the ramping and testing phases, within the 0% H_2_S group. We report confidence intervals of effect sizes and associated *p*-values (two-tailed, α=0.05), with full model summaries included in Supplementary Materials.

## Results

Aquatic surface respiration (ASR) rate in response to exogenous H_2_S was quantified by time spent at the surface at 21 and 28°C. The probability of surface respiration greatly increased with H_2_S level (Figure 1; Credible Intervals, CrI: 1.41–4.36) as well as with higher temperatures (CrI: 0.11–2.43). Our experimental design precluded hypoxia or H_2_S gradients; this suggests that ^ASR was driven by a reflexive respiratory drive rather than detection of reduced H^2^S near the^ surface. Because the respiratory response was reduced at lower temperatures, we examined whether fish would seek lower temperatures when challenged with H_2_S. Fish learned to control water temperature by swimming between the left (cold) or right (warm) chambers of the shuttlebox (e.g., highlighted individual traces in Figure 1B). Fish only exhibited overall side preferences with exogenous H_2_S (Figure 1 Bi, Ci; side preference in 0% H_2_S Confidence Interval, CI: −0.18–0.06; in 0.02% H_2_S CI: 0.21–0.48). Upon H_2_S exposure, fish rapidly selected reduced T_B_ (Figure 1 B ii, C ii; *p*<0.001; ΔT_B_ CI: 3.6–8.4°C), and exhibited reduced lower (*p*=0.002; ΔLET CI: 1.6–6.1) and upper (*p*=0.001; ΔUET CI: 1.7–6.7) escape temperatures.

Several fish entered the cold side and stopped shuttling altogether (Figure 1 Bii), which could mean that metabolic stress drives fish to escape and become trapped on the cold side. However, shuttling rates did not differ overall (*p*=0.60; CI: −0.23–0.40 min^−1^), and in fact swim velocity markedly increased after the introduction of gas (Figure 1 Ciii), which was sustained in the testing phase (*p*=0.007; CI: 0.21-0.68 cm s^−1^). Rather, side selection was dependent on variation in individual temperature preference (Figure 1 Di), despite acclimation together at 27°C for longer than a typical period of 10-12 days (e.g., [19,20]). The consistency of preferred T_B_ over the experiment duration in 0% H_2_S (e.g., individual and average traces, Figure 1 B) suggested to us that average T_B_ during the ramping phase could be used to gauge how temperatures are selected in the testing phase (Figure 1 B). In 0% H_2_S, a constant preferred temperature is indicated by a regression slope overlapping unity (Figure 1 D ii; slope 0.88, CI: 0.24–1.53) and reasonably high repeatability (*R*=0.54, CI: 0.13-0.78, *p*=0.006). The high correlation and conserved preference might be surprising given that fish are learning the paradigm during the ramping phase. The effect of learning instead appears to be in the shuttling rate, which is similarly repeatable (Figure 1 D iii, *log* shuttle rate *R*=0.51, CI: 0.08-0.77, *p*=0.01) but with slope less than unity (CI: 0.38-0.98; no significant effect of treatment, H_2_S *p*=0.85; *log* shuttling rate ✕ H_2_S: *p*=0.37). The low slope suggests that fish finetune behaviour to maintain preferred T_B_ with less effort. Given the repeatable T_B_, we examined how H_2_S exposure alters the thermal setpoint. If H_2_S causes a reduction in a fish’s innate T_B_ set point, we would expect to see an intercept difference alone between the ramping and test T_B_ relationship. Conversely, if H_2_S causes severe distress and an escape response so that fish try to achieve the lowest possible temperature regardless of their innate T_B_ set point, we would expect to observe a significant T_B_ ✕ H_2_S interaction. The intercept shift was not accompanied by an interaction (Figure 1 D ii; *p*<0.001; CI ΔT_B_: 4.1–8.3°C; T_B_ ✕ H_2_S: *p*=0.61), pointing to a reduction of T_B_ set point.

## Discussion

A central question in thermoregulatory physiology is the nature of the thermal setpoint and how it is adjusted [14,17]. Behavioural thermoregulation enables exploitation of the environment to select body temperatures that reflect the animal’s internal state. We found that individual adult zebrafish have temperature preferenda spanning at least 10°C (Figure 1), possibly due to factors such as juvenile growth conditions and standard metabolic rate [17] or health and age [20] (though no individuals were obviously ill, as judged by condition and robust escape response before testing). In response to the prevalent aquatic stressor H_2_S, fish temperature preference was reduced by a constant ~6°C though relative differences in preferenda were maintained (Figure 1 D ii). The preferences for lower temperatures with H_2_S exposure occurred despite higher overall activity (swim speed) but constant shuttling frequencies, which points to active defense of a new mean (e.g., exemplary individuals in Figure 1 B). Taken together, these observations point to a shift in innate thermal setpoint, similar to the widely conserved anapyrexic response observed in hypoxia [9,23,33]. In preliminary experiments testing the shuttlebox design, we examined thermoregulatory behaviour to 2% O_2_ which reliably elicits hypoxic responses [24]. In hypoxia, the change in body temperature was about half that observed with H_2_S (one-tailed *t*-test with unequal variances, *t*_11.4_=2.01, *p*=0.03, ΔT_B_ ~ −3.2°C), whereas swim speed decreased rather than increased (*t*_13.9_=1.85, *p*=0.04, Δ speed ~ 0.90; see also [24]). The blunter response to hypoxia alone is consistent with the emerging view that H_2_S is a key effector of the hypoxia response [4,25,34]. An important cellular candidate for responses to low oxygen are fishes’ neuroepithelial cells (NECs, [24]), which also contain H_2_S-producing enzymes [25]. Exogenous H_2_S greatly increases ventilatory rates and accentuates physiological responses to hypoxia [8], and partially rescues hypoxic ventilatory responses when NECs are inhibited [25]. Whether exogenous H_2_S is also sufficient to activate anapyrexia in terrestrial animals without some hypoxia is less clear [7,8], but it is promising that physiological similarities of mammalian carotid bodies and NECs include their responses to H_2_S [9,13,25]. A better understanding of the cellular mechanisms underlying behavioural responses in fish can therefore shed light more widely on the role and utility of H_2_S in functions such as artificial hibernation [7].

Hydrogen sulfide drives fish to seek alternative environments when possible, including through emersion [1,10] or refuge in habitats such as estuaries [3]. We found that fish also seek colder temperatures when exposed to H_2_S. The aquatic surface respiration rate was somewhat reduced when fish were held at 21°C rather than 28°C, temperatures that roughly coincide with the average thermal preference in each group (mean T_B_ 0% H_2_S: 27.3°C, 0.02% H_2_S: 21.3°C). We propose that environmental H_2_S can impact daily and seasonal habitat selection [1,20] by driving fish to cooler waters, which may contribute to intraspecific segregation and selection during repeated colonisations of H_2_S-replete habitats [10,12]. An important consideration is the value fish place in maintaining a habitat or defending a territory [21,22,26]: fish with higher temperature preference might be more resistant overall and less likely to abandon a current habitat in favour of searching for alternative environments. Whether this is ultimately advantageous will depend on the degree of environmental H_2_S and the combination of direct physiological and indirect impacts on microfauna and flora [3] that affect habitat suitability. Moreover, searching for cold refugia might in some cases be maladaptive because low temperatures can drive redox reactions that release H_2_S from mud [35]. Overall, we find that the capacity of H_2_S to alter behavioural thermal preferences in the absence of hypoxia [8] can contribute to its already complex environmental effects [1,3]. The potency of this effect appears to be due to its critical role in sensing and responding to oxygen levels, demonstrating that environmental hijacking of an endogenous gasotransmitter can profoundly affect animal behaviour.

## Supporting information

Supplementary Model Summaries

## Author contributions

JCS and GJT designed the shuttlebox; CDD, JCS, and GJT designed experiments; CDD and JCS performed experiments; DAS analysed results and prepared figures; DAS, CDD, and GJT wrote the manuscript; DAS and GJT approved the manuscript in its final form.

## Acknowledgements

We are especially grateful to Brock University’s Technical Services team for building the electronic shuttle box, and Viviana Cadena, Jacob Berman, and Qian Long for assistance with experiments, and to Miriam Richards for guidance with the behavioural assessment. This research was funded by Natural Sciences and Engineering Research Council of Canada (NSERC) Discovery Grants RGPIN-2014-05814 to GJT.

## Data availability

Data are available from the Dryad Digital Repository at: ##########

