## Supplementary Model Summaries for "Hydrogen sulfide exposure reduces thermal set point in zebrafish"

Model summaries for Skandalis et al.


### Model summaries for Skandalis et al.

###### 05 February 2020

- ANALYSIS OF HYDROGEN SULFIDE (0.02% H_2_S)
  - AQUATIC SURFACE RESPIRATION RATES BY TEMPERATURE AND H_2_S LEVEL
    - Generalised linear model of ASR ~ Temperature x H_2_S
  - POPULATION-AVERAGE BODY TEMPERATURE OVER TIME AND H_2_S
    - Generalised additive model of population fish temperature ~ Time (hrs) x H_2_S
  - AVERAGE TREATMENT EFFECTS
    - Fish temperature ~ H_2_S
    - Lower escape temperature ~ H_2_S
    - Upper escape temperature ~ H_2_S
    - log Shuttle rate ~ H_2_S
    - log Swim velocity ~ H_2_S
    - Time ratio ~ H_2_S
  - REPEATABILITY (0% H_2_S)
    - Fish temperature ramping->testing repeatability
    - Fish log shuttle rate ramping->testing repeatability
  - RESPONSES IN RAMPING AND TESTING PHASES
    - Fish temperature (testing) ~ Fish temperature (ramping) x H_2_S level
    - log Shuttle rate (testing) ~ log Shuttle rate (ramping) x H_2_S level
    - Lower escape temperature (testing) ~ Lower escape temperature (ramping) x H_2_S level
    - Upper escape temperature (testing) ~ Upper escape temperature (ramping) x H_2_S level
    - Swim velocity (testing) ~ Swim velocity (ramping) x H_2_S level
  - RELATIONSHIP OF FISH TEMPERATURE TO SWIM VARIABLES
    - Fish temperature ~ log Shuttle rate x H_2_S level
    - Fish temperature ~ Swim velocity x H_2_S level
    - Fish temperature ~ Time ratio x H_2_S level
- ANALYSIS OF HYPOXIA (2% O_2_)
  - AVERAGE TREATMENT EFFECTS
    - Fish temperature ~ O_2_
    - Lower escape temperature ~ O_2_
    - Upper escape temperature ~ O_2_
    - Swim velocity ~ O_2_
    - log Shuttle rate ~ O_2_
    - Side preference ~ O_2_

### ANALYSIS OF HYDROGEN SULFIDE (0.02% H_2_S)

Aquatic surface respiration data collected by Glenn J. Tattersall

Shuttlebox data collected by Cheryl D. Dobell

Analysis performed by Dimitri A. Skandalis

#### AQUATIC SURFACE RESPIRATION RATES BY TEMPERATURE AND H_2_S LEVEL

##### Generalised linear model of ASR ~ Temperature x H_2_S

*ASR* is the number of frames in which the fish is near the surface. *n* is the total number of frames.

```
##  Family: binomial 
##   Links: mu = logit 
## Formula: ASR | trials(n) ~ Temperature * H2S + (1 | ID/H2S/Temperature) 
##    Data: dat_asr_summary (Number of observations: 26) 
## Samples: 4 chains, each with iter = 10000; warmup = 5000; thin = 1;
##          total post-warmup samples = 20000
## 
## Group-Level Effects: 
## ~ID (Number of levels: 8) 
##               Estimate Est.Error l-95% CI u-95% CI Rhat Bulk_ESS Tail_ESS
## sd(Intercept)     0.78      0.52     0.05     2.03 1.00     8171    10504
## 
## ~ID:H2S (Number of levels: 14) 
##               Estimate Est.Error l-95% CI u-95% CI Rhat Bulk_ESS Tail_ESS
## sd(Intercept)     0.67      0.41     0.04     1.60 1.00     6286     9296
## 
## ~ID:H2S:Temperature (Number of levels: 26) 
##               Estimate Est.Error l-95% CI u-95% CI Rhat Bulk_ESS Tail_ESS
## sd(Intercept)     0.97      0.24     0.60     1.55 1.00     8103    11670
## 
## Population-Level Effects: 
##                       Estimate Est.Error l-95% CI u-95% CI Rhat Bulk_ESS Tail_ESS
## Intercept                -4.22      0.64    -5.51    -2.97 1.00    14684    12257
## Temperature28             1.22      0.59     0.12     2.43 1.00    13100    10906
## H2S0.02                   2.84      0.75     1.40     4.39 1.00    13457    12289
## Temperature28:H2S0.02    -1.20      0.83    -2.91     0.38 1.00    12168    11072
## 
## Samples were drawn using sampling(NUTS). For each parameter, Bulk_ESS
## and Tail_ESS are effective sample size measures, and Rhat is the potential
## scale reduction factor on split chains (at convergence, Rhat = 1).
```

#### POPULATION-AVERAGE BODY TEMPERATURE OVER TIME AND H_2_S

##### Generalised additive model of population fish temperature ~ Time (hrs) x H_2_S

*Treat\_level.L* is the main effect of H_2_S exposure (0% is reference). *s(t.hr)* is the reference smooth. *s(t.hr):Treat\_levelH2S\_0.02%* is the smooth of differences from the reference.

```
## 
## Family: gaussian 
## Link function: identity 
## 
## Formula:
## Temp.fish ~ s(t.hr, bs = "gp") + s(t.hr, by = Treat_level, bs = "gp") + 
##     Treat_level
## 
## Parametric coefficients:
##                Estimate Std. Error t value Pr(>|t|)    
## (Intercept)   25.731613   0.005582  4609.5   <2e-16 ***
## Treat_level.L -2.118710   0.007895  -268.4   <2e-16 ***
## ---
## Signif. codes:  0 '***' 0.001 '**' 0.01 '*' 0.05 '.' 0.1 ' ' 1
## 
## Approximate significance of smooth terms:
##                                edf Ref.df       F p-value    
## s(t.hr)                      11.00     11   166.7  <2e-16 ***
## s(t.hr):Treat_levelH2S_0.02% 10.99     11 12522.3  <2e-16 ***
## ---
## Signif. codes:  0 '***' 0.001 '**' 0.01 '*' 0.05 '.' 0.1 ' ' 1
## 
## R-sq.(adj) =  0.406   Deviance explained = 40.6%
## GCV = 12.157  Scale est. = 12.157    n = 403624
```

#### AVERAGE TREATMENT EFFECTS

##### Fish temperature ~ H_2_S

```
## 
## Call:
## lm(formula = Temp.fish ~ Treat_level, data = dat_CDD_testphase)
## 
## Residuals:
##     Min      1Q  Median      3Q     Max 
## -6.6897 -2.5802  0.2867  1.8745  7.4603 
## 
## Coefficients:
##                      Estimate Std. Error t value Pr(>|t|)    
## (Intercept)           27.2977     0.7916  34.485  < 2e-16 ***
## Treat_levelH2S_0.02%  -6.0016     1.1678  -5.139 1.05e-05 ***
## ---
## Signif. codes:  0 '***' 0.001 '**' 0.01 '*' 0.05 '.' 0.1 ' ' 1
## 
## Residual standard error: 3.54 on 35 degrees of freedom
## Multiple R-squared:  0.4301, Adjusted R-squared:  0.4138 
## F-statistic: 26.41 on 1 and 35 DF,  p-value: 1.053e-05
```

Confidence intervals:

```
##                          2.5 %   97.5 %
## (Intercept)          25.690658 28.90468
## Treat_levelH2S_0.02% -8.372442 -3.63084
```

##### Lower escape temperature ~ H_2_S

```
## 
## Call:
## lm(formula = LET ~ Treat_level, data = dat_CDD_testphase)
## 
## Residuals:
##     Min      1Q  Median      3Q     Max 
## -7.2700 -2.4587  0.8313  2.2224  6.1822 
## 
## Coefficients:
##                      Estimate Std. Error t value Pr(>|t|)    
## (Intercept)           25.9272     0.7545  34.361  < 2e-16 ***
## Treat_levelH2S_0.02%  -3.8111     1.1132  -3.424  0.00159 ** 
## ---
## Signif. codes:  0 '***' 0.001 '**' 0.01 '*' 0.05 '.' 0.1 ' ' 1
## 
## Residual standard error: 3.374 on 35 degrees of freedom
## Multiple R-squared:  0.2509, Adjusted R-squared:  0.2295 
## F-statistic: 11.72 on 1 and 35 DF,  p-value: 0.001591
```

Confidence intervals:

```
##                          2.5 %    97.5 %
## (Intercept)          24.395354 27.458972
## Treat_levelH2S_0.02% -6.070921 -1.551207
```

##### Upper escape temperature ~ H_2_S

```
## 
## Call:
## lm(formula = UET ~ Treat_level, data = dat_CDD_testphase)
## 
## Residuals:
##     Min      1Q  Median      3Q     Max 
## -8.3378 -2.3257  0.7727  2.1422  6.4464 
## 
## Coefficients:
##                      Estimate Std. Error t value Pr(>|t|)    
## (Intercept)           29.3092     0.8228  35.622  < 2e-16 ***
## Treat_levelH2S_0.02%  -4.2056     1.2138  -3.465  0.00142 ** 
## ---
## Signif. codes:  0 '***' 0.001 '**' 0.01 '*' 0.05 '.' 0.1 ' ' 1
## 
## Residual standard error: 3.68 on 35 degrees of freedom
## Multiple R-squared:  0.2554, Adjusted R-squared:  0.2341 
## F-statistic:    12 on 1 and 35 DF,  p-value: 0.001421
```

Confidence intervals:

```
##                          2.5 %    97.5 %
## (Intercept)          27.638914 30.979563
## Treat_levelH2S_0.02% -6.669834 -1.741419
```

##### log Shuttle rate ~ H_2_S

```
## 
## Call:
## lm(formula = log_Shuttle_rt ~ Treat_level, data = dat_CDD_testphase)
## 
## Residuals:
##      Min       1Q   Median       3Q      Max 
## -0.89466 -0.36173 -0.00825  0.32151  0.84276 
## 
## Coefficients:
##                      Estimate Std. Error t value Pr(>|t|)    
## (Intercept)           1.62893    0.10541  15.454   <2e-16 ***
## Treat_levelH2S_0.02%  0.08226    0.15550   0.529      0.6    
## ---
## Signif. codes:  0 '***' 0.001 '**' 0.01 '*' 0.05 '.' 0.1 ' ' 1
## 
## Residual standard error: 0.4714 on 35 degrees of freedom
## Multiple R-squared:  0.007931,   Adjusted R-squared:  -0.02041 
## F-statistic: 0.2798 on 1 and 35 DF,  p-value: 0.6002
```

Confidence intervals:

```
##                           2.5 %   97.5 %
## (Intercept)           1.4149457 1.842918
## Treat_levelH2S_0.02% -0.2334347 0.397947
```

##### log Swim velocity ~ H_2_S

```
## 
## Call:
## lm(formula = log(Velocity) ~ Treat_level, data = dat_CDD_testphase)
## 
## Residuals:
##     Min      1Q  Median      3Q     Max 
## -0.8365 -0.2284 -0.1433  0.2824  1.4661 
## 
## Coefficients:
##                      Estimate Std. Error t value Pr(>|t|)    
## (Intercept)           -1.4661     0.1167  -12.57 1.56e-14 ***
## Treat_levelH2S_0.02%   0.4905     0.1721    2.85  0.00727 ** 
## ---
## Signif. codes:  0 '***' 0.001 '**' 0.01 '*' 0.05 '.' 0.1 ' ' 1
## 
## Residual standard error: 0.5217 on 35 degrees of freedom
## Multiple R-squared:  0.1884, Adjusted R-squared:  0.1652 
## F-statistic: 8.124 on 1 and 35 DF,  p-value: 0.007274
```

Confidence intervals (on metric scale):

```
##     2.5 %    97.5 % 
## 0.2097651 0.6774714
```

##### Time ratio ~ H_2_S

```
## 
## Call:
## lm(formula = Time_ratio ~ Treat_level - 1, data = dat_CDD_testphase)
## 
## Residuals:
##      Min       1Q   Median       3Q      Max 
## -0.55589 -0.10930  0.00114  0.13199  0.84458 
## 
## Coefficients:
##                      Estimate Std. Error t value Pr(>|t|)    
## Treat_levelCTL_0%     0.05667    0.06216   0.912    0.368    
## Treat_levelH2S_0.02% -0.34491    0.06742  -5.116 1.13e-05 ***
## ---
## Signif. codes:  0 '***' 0.001 '**' 0.01 '*' 0.05 '.' 0.1 ' ' 1
## 
## Residual standard error: 0.278 on 35 degrees of freedom
## Multiple R-squared:  0.4355, Adjusted R-squared:  0.4033 
## F-statistic:  13.5 on 2 and 35 DF,  p-value: 4.508e-05
```

Confidence intervals:

```
##                            2.5 %     97.5 %
## Treat_levelCTL_0%    -0.06952303  0.1828530
## Treat_levelH2S_0.02% -0.48178175 -0.2080415
```

#### REPEATABILITY (0% H_2_S)

##### Fish temperature ramping->testing repeatability

```
## 
## Repeatability estimation using the lmm method
## 
## Call = rptR::rptGaussian(formula = Temp.fish ~ (1 | fishID), grname = "fishID", data = dat_icc, parallel = TRUE, ncores = 2, adjusted = TRUE)
## 
## Data: 40 observations
## ----------------------------------------
## 
## fishID (20 groups)
## 
## Repeatability estimation overview: 
##       R     SE   2.5%  97.5% P_permut  LRT_P
##   0.539  0.158   0.13  0.771       NA  0.006
## 
## Bootstrapping and Permutation test: 
##             N   Mean Median   2.5%  97.5%
## boot     1000  0.519  0.534   0.13  0.771
## permut      1     NA     NA     NA     NA
## 
## Likelihood ratio test: 
## logLik full model = -96.16956
## logLik red. model = -99.33616
## D  = 6.33, df = 1, P = 0.00592
## 
## ----------------------------------------
```

##### Fish log shuttle rate ramping->testing repeatability

```
## 
## Repeatability estimation using the lmm method
## 
## Call = rptR::rptGaussian(formula = log_Shuttle_rt ~ (1 | fishID), grname = "fishID", data = dat_icc, parallel = TRUE, ncores = 2, adjusted = TRUE)
## 
## Data: 40 observations
## ----------------------------------------
## 
## fishID (20 groups)
## 
## Repeatability estimation overview: 
##       R     SE   2.5%  97.5% P_permut  LRT_P
##   0.508  0.172  0.111  0.775       NA   0.01
## 
## Bootstrapping and Permutation test: 
##             N   Mean Median   2.5%  97.5%
## boot     1000  0.499  0.512  0.111  0.775
## permut      1     NA     NA     NA     NA
## 
## Likelihood ratio test: 
## logLik full model = -32.38444
## logLik red. model = -35.11355
## D  = 5.46, df = 1, P = 0.00974
## 
## ----------------------------------------
```

#### RESPONSES IN RAMPING AND TESTING PHASES

##### Fish temperature (testing) ~ Fish temperature (ramping) x H_2_S level

```
## 
## Call:
## lm(formula = Temp.fish_Test ~ Temp.fish_Ramping * Treat_level, 
##     data = dat_period)
## 
## Residuals:
##     Min      1Q  Median      3Q     Max 
## -7.4145 -2.6392 -0.1962  1.7412  7.2704 
## 
## Coefficients:
##                                        Estimate Std. Error t value Pr(>|t|)    
## (Intercept)                              2.8459     0.7033   4.047 0.000295 ***
## Temp.fish_Ramping                        0.8832     0.3155   2.799 0.008488 ** 
## Treat_levelH2S_0.02%                    -6.1661     1.0377  -5.942 1.14e-06 ***
## Temp.fish_Ramping:Treat_levelH2S_0.02%  -0.2362     0.4646  -0.508 0.614562    
## ---
## Signif. codes:  0 '***' 0.001 '**' 0.01 '*' 0.05 '.' 0.1 ' ' 1
## 
## Residual standard error: 3.142 on 33 degrees of freedom
## Multiple R-squared:  0.5767, Adjusted R-squared:  0.5383 
## F-statistic: 14.99 on 3 and 33 DF,  p-value: 2.508e-06
```

##### log Shuttle rate (testing) ~ log Shuttle rate (ramping) x H_2_S level

```
## 
## Call:
## lm(formula = log_Shuttle_rt_Test ~ log_Shuttle_rt_Ramping * Treat_level, 
##     data = dat_period)
## 
## Residuals:
##      Min       1Q   Median       3Q      Max 
## -0.86440 -0.27410  0.03254  0.27154  0.70360 
## 
## Coefficients:
##                                             Estimate Std. Error t value Pr(>|t|)    
## (Intercept)                                  0.02243    0.08278   0.271    0.788    
## log_Shuttle_rt_Ramping                       0.67825    0.14773   4.591 6.12e-05 ***
## Treat_levelH2S_0.02%                        -0.02381    0.12336  -0.193    0.848    
## log_Shuttle_rt_Ramping:Treat_levelH2S_0.02% -0.23932    0.26150  -0.915    0.367    
## ---
## Signif. codes:  0 '***' 0.001 '**' 0.01 '*' 0.05 '.' 0.1 ' ' 1
## 
## Residual standard error: 0.3655 on 33 degrees of freedom
## Multiple R-squared:  0.4376, Adjusted R-squared:  0.3865 
## F-statistic: 8.561 on 3 and 33 DF,  p-value: 0.0002409
```

##### Lower escape temperature (testing) ~ Lower escape temperature (ramping) x H_2_S level

```
## 
## Call:
## lm(formula = LET_Test ~ LET_Ramping * Treat_level, data = dat_period)
## 
## Residuals:
##     Min      1Q  Median      3Q     Max 
## -8.3965 -2.4839  0.7926  2.1082  6.4492 
## 
## Coefficients:
##                                  Estimate Std. Error t value Pr(>|t|)    
## (Intercept)                        1.7671     0.7104   2.487 0.018096 *  
## LET_Ramping                        0.7821     0.3389   2.308 0.027417 *  
## Treat_levelH2S_0.02%              -3.8367     1.0481  -3.661 0.000872 ***
## LET_Ramping:Treat_levelH2S_0.02%  -0.3882     0.4985  -0.779 0.441683    
## ---
## Signif. codes:  0 '***' 0.001 '**' 0.01 '*' 0.05 '.' 0.1 ' ' 1
## 
## Residual standard error: 3.177 on 33 degrees of freedom
## Multiple R-squared:  0.3739, Adjusted R-squared:  0.317 
## F-statistic:  6.57 on 3 and 33 DF,  p-value: 0.001321
```

##### Upper escape temperature (testing) ~ Upper escape temperature (ramping) x H_2_S level

```
## 
## Call:
## lm(formula = UET_Test ~ UET_Ramping * Treat_level, data = dat_period)
## 
## Residuals:
##     Min      1Q  Median      3Q     Max 
## -9.0588 -2.4412  0.4711  1.8661  6.6878 
## 
## Coefficients:
##                                  Estimate Std. Error t value Pr(>|t|)    
## (Intercept)                        1.9681     0.7999   2.460 0.019278 *  
## UET_Ramping                        0.5993     0.3383   1.771 0.085721 .  
## Treat_levelH2S_0.02%              -4.2691     1.1802  -3.617 0.000983 ***
## UET_Ramping:Treat_levelH2S_0.02%  -0.2044     0.5332  -0.383 0.703957    
## ---
## Signif. codes:  0 '***' 0.001 '**' 0.01 '*' 0.05 '.' 0.1 ' ' 1
## 
## Residual standard error: 3.576 on 33 degrees of freedom
## Multiple R-squared:  0.3369, Adjusted R-squared:  0.2766 
## F-statistic: 5.589 on 3 and 33 DF,  p-value: 0.003261
```

##### Swim velocity (testing) ~ Swim velocity (ramping) x H_2_S level

```
## 
## Call:
## lm(formula = Velocity_Test ~ Velocity_Ramping * Treat_level, 
##     data = dat_period)
## 
## Residuals:
##      Min       1Q   Median       3Q      Max 
## -0.31011 -0.08899 -0.01745  0.06981  0.67637 
## 
## Coefficients:
##                                       Estimate Std. Error t value Pr(>|t|)  
## (Intercept)                           -0.04535    0.03929  -1.154   0.2567  
## Velocity_Ramping                       0.13320    0.07422   1.795   0.0819 .
## Treat_levelH2S_0.02%                   0.10358    0.05797   1.787   0.0831 .
## Velocity_Ramping:Treat_levelH2S_0.02% -0.11541    0.10262  -1.125   0.2689  
## ---
## Signif. codes:  0 '***' 0.001 '**' 0.01 '*' 0.05 '.' 0.1 ' ' 1
## 
## Residual standard error: 0.1753 on 33 degrees of freedom
## Multiple R-squared:  0.1718, Adjusted R-squared:  0.09649 
## F-statistic: 2.282 on 3 and 33 DF,  p-value: 0.0974
```

#### RELATIONSHIP OF FISH TEMPERATURE TO SWIM VARIABLES

##### Fish temperature ~ log Shuttle rate x H_2_S level

```
## 
## Call:
## lm(formula = Temp.fish_Test ~ log_Shuttle_rt_Test * Treat_level, 
##     data = dat_period, plot = FALSE)
## 
## Residuals:
##     Min      1Q  Median      3Q     Max 
## -6.1966 -2.1548 -0.1841  2.0549  8.0796 
## 
## Coefficients:
##                                          Estimate Std. Error t value Pr(>|t|)    
## (Intercept)                                2.7898     0.8128   3.432  0.00163 ** 
## log_Shuttle_rt_Test                        0.8533     1.5117   0.564  0.57627    
## Treat_levelH2S_0.02%                      -6.0573     1.2029  -5.036 1.66e-05 ***
## log_Shuttle_rt_Test:Treat_levelH2S_0.02%  -0.3261     2.9631  -0.110  0.91304    
## ---
## Signif. codes:  0 '***' 0.001 '**' 0.01 '*' 0.05 '.' 0.1 ' ' 1
## 
## Residual standard error: 3.626 on 33 degrees of freedom
## Multiple R-squared:  0.4362, Adjusted R-squared:  0.385 
## F-statistic: 8.512 on 3 and 33 DF,  p-value: 0.0002506
```

##### Fish temperature ~ Swim velocity x H_2_S level

```
## 
## Call:
## lm(formula = Temp.fish_Test ~ Velocity_Test * Treat_level, data = dat_period, 
##     plot = FALSE)
## 
## Residuals:
##     Min      1Q  Median      3Q     Max 
## -6.5719 -2.4624  0.3197  1.8049  7.3732 
## 
## Coefficients:
##                                    Estimate Std. Error t value Pr(>|t|)    
## (Intercept)                          2.7062     0.8377   3.231   0.0028 ** 
## Velocity_Test                       -1.0244     3.9146  -0.262   0.7952    
## Treat_levelH2S_0.02%                -5.9709     1.2911  -4.625 5.55e-05 ***
## Velocity_Test:Treat_levelH2S_0.02%   1.3743     8.2767   0.166   0.8691    
## ---
## Signif. codes:  0 '***' 0.001 '**' 0.01 '*' 0.05 '.' 0.1 ' ' 1
## 
## Residual standard error: 3.642 on 33 degrees of freedom
## Multiple R-squared:  0.4313, Adjusted R-squared:  0.3796 
## F-statistic: 8.342 on 3 and 33 DF,  p-value: 0.0002881
```

##### Fish temperature ~ Time ratio x H_2_S level

```
## 
## Call:
## lm(formula = Temp.fish_Test ~ Time_ratio_Test, data = dat_period, 
##     plot = FALSE)
## 
## Residuals:
##     Min      1Q  Median      3Q     Max 
## -6.9229 -1.5536 -0.1451  2.0979  4.1605 
## 
## Coefficients:
##                   Estimate Std. Error t value Pr(>|t|)    
## (Intercept)     -1.851e-15  4.599e-01   0.000        1    
## Time_ratio_Test  1.088e+01  1.367e+00   7.959 2.31e-09 ***
## ---
## Signif. codes:  0 '***' 0.001 '**' 0.01 '*' 0.05 '.' 0.1 ' ' 1
## 
## Residual standard error: 2.797 on 35 degrees of freedom
## Multiple R-squared:  0.6441, Adjusted R-squared:  0.634 
## F-statistic: 63.35 on 1 and 35 DF,  p-value: 2.311e-09
```

### ANALYSIS OF HYPOXIA (2% O_2_)

Shuttlebox data collected by Joshua C. Shaw

Analysis performed by Dimitri A. Skandalis

#### AVERAGE TREATMENT EFFECTS

##### Fish temperature ~ O_2_

```
## 
##  Welch Two Sample t-test
## 
## data:  Temp.fish by Treat_level
## t = 2.0056, df = 11.384, p-value = 0.03463
## alternative hypothesis: true difference in means is greater than 0
## 95 percent confidence interval:
##  0.3388087       Inf
## sample estimates:
## mean in group NOX_21%  mean in group HPX_2% 
##            -0.4819944            -3.6385575
```

##### Lower escape temperature ~ O_2_

```
## 
##  Welch Two Sample t-test
## 
## data:  LET by Treat_level
## t = 2.0246, df = 13.882, p-value = 0.03129
## alternative hypothesis: true difference in means is greater than 0
## 95 percent confidence interval:
##  0.2355009       Inf
## sample estimates:
## mean in group NOX_21%  mean in group HPX_2% 
##             -1.787126             -3.605332
```

##### Upper escape temperature ~ O_2_

```
## 
##  Welch Two Sample t-test
## 
## data:  UET by Treat_level
## t = 1.8713, df = 13.802, p-value = 0.04133
## alternative hypothesis: true difference in means is greater than 0
## 95 percent confidence interval:
##  0.09558922        Inf
## sample estimates:
## mean in group NOX_21%  mean in group HPX_2% 
##             1.2600558            -0.3938188
```

##### Swim velocity ~ O_2_

```
## 
##  Welch Two Sample t-test
## 
## data:  Velocity by Treat_level
## t = 1.8521, df = 13.895, p-value = 0.04269
## alternative hypothesis: true difference in means is greater than 0
## 95 percent confidence interval:
##  0.04384515        Inf
## sample estimates:
## mean in group NOX_21%  mean in group HPX_2% 
##            -0.4857143            -1.3900000
```

##### log Shuttle rate ~ O_2_

```
## 
##  Welch Two Sample t-test
## 
## data:  log_Shuttle_rt by Treat_level
## t = 1.543, df = 14.585, p-value = 0.07212
## alternative hypothesis: true difference in means is greater than 0
## 95 percent confidence interval:
##  -0.03660821         Inf
## sample estimates:
## mean in group NOX_21%  mean in group HPX_2% 
##           -0.03416448           -0.29898407
```

##### Side preference ~ O_2_

```
## 
##  Welch Two Sample t-test
## 
## data:  Time_ratio by Treat_level
## t = 1.091, df = 9.529, p-value = 0.1511
## alternative hypothesis: true difference in means is greater than 0
## 95 percent confidence interval:
##  -0.06543593         Inf
## sample estimates:
## mean in group NOX_21%  mean in group HPX_2% 
##            0.03609128           -0.06161372
```
